# Scalable full-transcript coverage single cell RNA sequencing with Smart-seq3xpress

**DOI:** 10.1101/2021.07.10.451889

**Authors:** Michael Hagemann-Jensen, Christoph Ziegenhain, Rickard Sandberg

## Abstract

Plate-based single-cell RNA-sequencing methods with full-transcript coverage typically excel at sensitivity but are more costly and time-consuming. Here, we miniaturized and streamlined the Smart-seq3 protocol for drastically reduced cost and increased throughput. Applying Smart-seq3xpress to 16,349 human peripheral blood mononuclear cells revealed a highly granular atlas complete with both common and rare cell types whose identification previously relied on additional protein measurements or the integration with a reference atlas.

---

Large numbers of single-cell RNA-sequencing methods have been described in recent years and their respective strengths and weaknesses have been well characterised^1,2^. However, researchers are still confronted with striking a compromise between methods with high cellular throughput (i.e. droplet or combinatorial indexing methods) or methods with high sensitivity and full-length transcript coverage (i.e. plate-based methods). Based on Smart-seq3^3^, the method currently offering the highest information content per profiled cell, we systematically evaluated the feasibility to reduce volumes, reagents, and experimental steps, without sacrificing data quality. This resulted in Smart-seq3xpress, a scalable nanoliter implementation of Smart-seq3 with a cost of 0.25 EUR per cell - comparable to generating 7,000 cells with 10x Genomics. Throughput is only limited by the available equipment (e.g. PCR instruments) and crucially, sequencing-ready libraries can be generated within a single workday.

We hypothesized that the Smart-seq3 chemistry would work in significantly lower volumes without sacrificing quality if sufficiently protected from evaporation, e.g. when covered in a inert hydrophobic substance (“overlay”)^4^ (**Figure 1a**). Using accurate non-contact nanoliter dispensers, we scaled the reaction volumes of the lysis, reverse transcription and pre-amplification PCR steps down to 1:2, 1:5 and 1:10 of the established volumes and tested these conditions on K562 and HEK293FT cells (**Figure 1b**). For the lowest volumes, cells were sorted by FACS into 300 nL of lysis buffer covered with 3 μL of VaporLock, with subsequent additions of 100 nL for reverse transcription and 600 nL for PCR. After shallow sequencing, alignment and error correction of reads, we observed similar numbers of detected genes and molecules per cell at a sequencing depth of 100,000 reads per cell (**Figure 1c-d, Supplementary Figure 1**), confirming that reaction scaling is possible without compromising data quality. Further reduction of reaction volumes beyond 1:10 were also possible (data not shown), although not further pursued as cost savings are diminishing and reactions may become vulnerable to variations in cell sorting fluid (~5 nL). We hypothesized that the overlays would both protect the low reaction volumes from evaporation and provide a “landing cushion” for the FACS sorted cells. Indeed, many overlays with varying chemical properties could be utilized with low-volume Smart-seq3 (**Figure 1f**), including Silicone oils with high viscosities and hydrocarbons with higher freeze points. As expected, overlays did not interfere with the cDNA synthesis reaction when tested in larger volumes (**Supplementary Figure 2a**). Performing low-volume Smart-seq3 without an overlay resulted in significant losses (**Figure 1f**), which could explain why earlier efforts to miniaturize plate-based scRNA-seq methods that did not utilize an inert overlay resulted in significantly decreased complexity^5–7^. Next, we investigated whether the time- and plastics-consuming cDNA clean-up step could simply be omitted by instead diluting the cDNA obtained in lower volumes. At equal sequencing depths, single-cell libraries generated with cDNA dilution and clean-up had no detectable differences (p=0.89 and p=0.71, respectively, *t*-test; **Figure 1e**). The significantly lowered reaction volumes enabled better control of downstream reaction conditions, and we therefore investigated the relationship between RT and preamplification PCR volumes. We found that dilution of the RT reaction is beneficial for optimal PCR performance, and we kept the established 1.5x ratio of PCR to RT volume (**Supplementary Figure 3**), and that PCR extension times could be reduced from 6 to 4 minutes, whereas further shortenings resulted in complexity losses, especially for longer transcripts (**Supplementary Figure 4a**).

**Figure 1.**
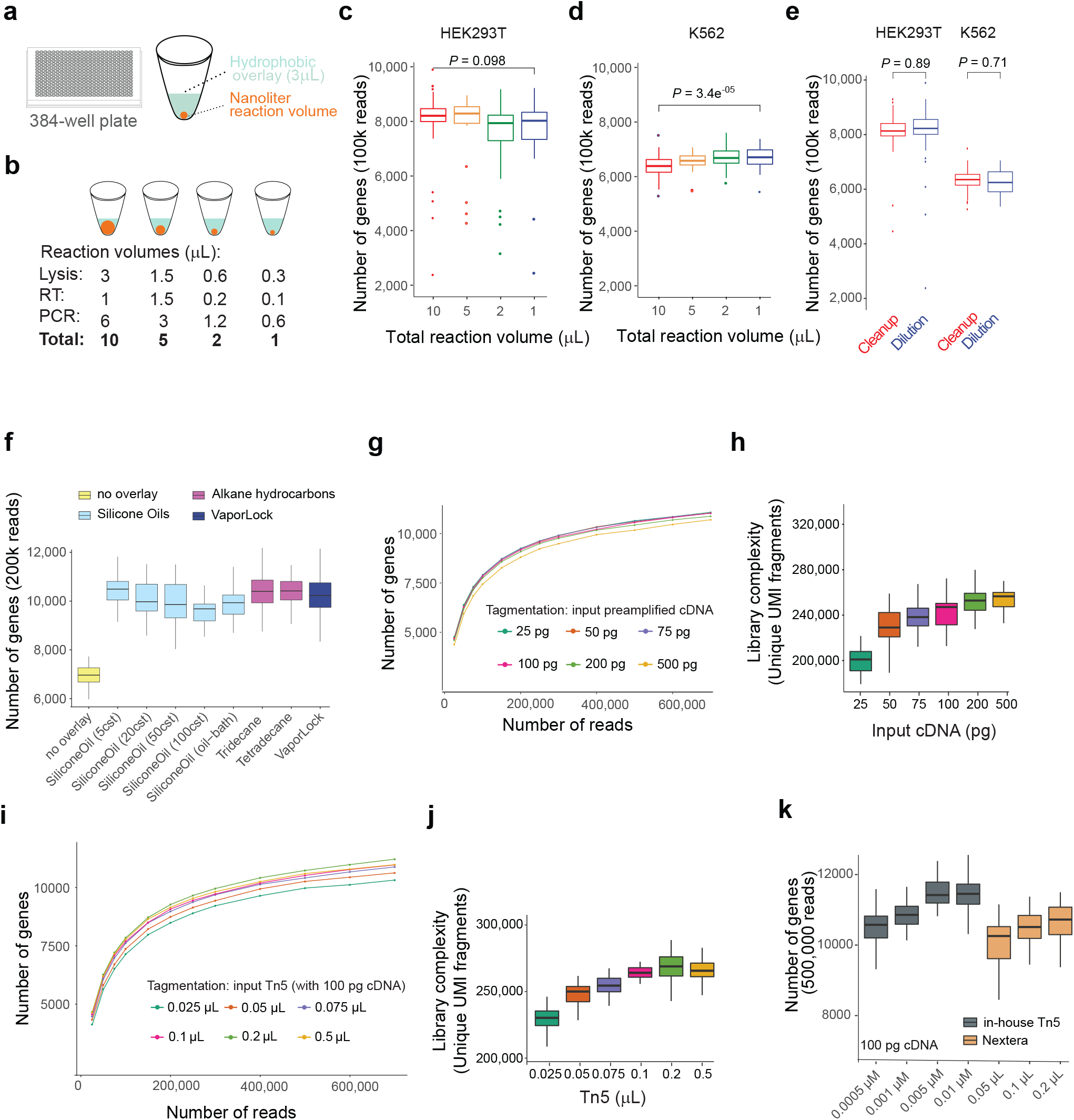
Optimization of a nanoliter Smart-seq3 workflow. (**a**) Schematic of nanoliter cDNA synthesis reactions performed in wells of 384-well PCR plates with 3 μL hydrophobic overlay. (**b**) Illustration of the volume reduction experiment with the exact lysis, reverse transcription (RT) and PCR volumes used. (**c**) Reduction of reaction volumes in single HEK293FT cells. Shown are the number of reads detected per cell at each reaction volume when sampling 100,000 sequencing reads (n = 100, 19, 32, 28 cells, respectively). P-value represents a t test between the 10 μL and 1 μL conditions. (**d**) Reduction of reaction volumes in single K562 cells. Shown are the number of reads detected per cell at each reaction volume when sampling 100,000 sequencing reads (n = 63, 39, 55, 53 cells, respectively). P-value represents a t test between the 10 μL and 1 μL conditions. (**e**) Replacement of the bead-based cDNA cleanup by dilution in single HEK293FT (n = 58, 52, respectively) and K562 (n = 57, 38, respectively) cells. Shown are the number of reads detected per cell for each condition at 100,000 reads with a p-value for a t-test within cell types. (**f**) Influence of hydrophobic overlays on miniaturized cDNA synthesis (1 μL total volume). For each compound, boxes depict the number of Genes detected from single HEK293FT cells (n = 17, 34, 39, 28, 25, 24, 28, 38, 70, respectively) when sampling 200,000 sequencing reads per cell. (**g**) Tagmentation complexity using 0.1 μL ATM Tn5 enzyme per single HEK293FT cell in relation to the input cDNA. For each dot, the median number of detected Genes is calculated from the indicated number of raw sequencing reads and plotted from at least n>=5 cells. (**h**) For varying amounts of cDNA input (see (g)), tagmentation complexity was summarized as unique aligned and gene-assigned UMI-containing read-pairs per 400,000 raw reads per HEK293FT cell (n = 49, 51, 51, 50, 51, 44). (**i**) Tagmentation complexity using 100 pg cDNA per single HEK293FT cell in relation to the enzyme amount (ATM Tn5). For each dot, the median number of detected Genes is calculated from the indicated number of raw sequencing reads and plotted from at least n>=5 cells. (**j**) For varying amounts of Tn5 enzyme (see (i)), tagmentation complexity was summarized as unique aligned and gene-assigned UMI-containing read-pairs per 400,000 raw reads per HEK293FT cell (n = 53, 46, 56, 57, 52, 50, respectively). (**k**) The use of in-house Tn5 relative to performance of ATM Tn5 was compared in HEK293FT cells. Each boxplot shows the number of Genes detected at 500,000 raw reads (n = 182, 52, 24, 226, 41, 53, 50, respectively).

A major cost in plate-based scRNA-seq is tagmentation which needs to be performed individually on each cell, and although reducing the amounts of commercial Tn5 has been suggested to cut reaction costs^8,9^, it is currently unclear to what degree Tn5 can be reduced without losing library complexity. We first investigated how variations of the input amounts of cDNA influence library complexity, which revealed that 20-fold variations in cDNA could be tolerated with minor effects on gene detection (**Figure 1g**) and unique fragments sequenced (**Figure 1h**). We further confirmed the notion that complexity is mainly a product of absolute Tn5 and cDNA amounts with little impact from varying reaction volumes (**Supplementary Figure 5**). The slightly reduced complexity at the highest cDNA amounts (**Figure 1g**) likely resulted from insufficient tagmentation due to the limited Tn5 amounts used. Similar results were observed when varying the Tn5 amounts on a fixed amount of cDNA input (**Figure 1i,j**) and high-quality libraries were obtained using both commercial and in-house^10^ Tn5 (**Figure 1k**). Altogether, these results demonstrated that tagmentation reactions were robust over large variations in cDNA input and Tn5 amounts, and that significant cost-savings can be made in reducing Tn5 amounts with only minimal impact on library complexity (**Figure 1h,j**).

Having demonstrated robust tagmentation over large ranges of cDNA, Tn5 and reaction volumes, we realized an opportunity to exclude several time-consuming and expensive experimental steps, including excessive cDNA preamplification, concentration measurements, bioanalyzer QC traces and cDNA amount normalization (**Figure 2a**). Instead, cDNA products after preamplification at fewer cycles could directly be tagmented without the above-mentioned steps – this strategy we termed Smart-seq3xpress. To explore this potential, we first generated libraries with low-volume RT (300 nL lysis volume) from HEK293FT cells and using a range of PCR preamplification cycles (10 to 20), that strikingly, revealed very similar gene detection (**Figure 2b**) without any need for additional enzymatic reaction clean-ups (**Supplementary Figure 8a).** However, the resulting libraries were heavily biased towards 5’ reads that contain the UMI at the expense of the internal reads important for full-length scRNA-seq as for the established Smart-seq3 protocol^3^, thus the resulting libraries had diminished the full-transcript coverage. This resulted from inefficient tagmentation, evident in the inability to modulate the ratio of UMI-containing and internal reads with increasing Tn5 amounts (**Figure 2c**). Specifically, the high salt concentration in the KAPA PCR buffer likely resulted in template blocking during tagmentation (https://emea.illumina.com/content/dam/illumina-marketing/documents/products/technotes/nextera-xt-troubleshooting-technical-note.pdf). However, five other PCR polymerases were compatible with direct tagmentation in terms of library quality and complexity (**Supplementary Figure 6a,b**). Importantly, tagmentation of SeqAmp and PlatinumII amplified cDNA had significantly lowered fraction of 5’ UMI reads indicative of improved tagmentation compatibility (**Supplementary Figure 6c**). Additional experiments identified SeqAmp as the preferred PCR polymerase, as it consistently performed well with improved gene and molecule detection over a range of template switching oligonucleotide (TSO) and PCR primer amounts (**Supplementary Figure 4b** and **6d**). Moreover, we used molecular spikes^11^ to directly assess the accuracy of RNA counting. Whereas PlatinumII had higher error-rates which resulted in inflated RNA counts (**Supplementary Figure 7**), SeqAmp with 2 μM TSO and 1 μM of each PCR primer had high sensitivity and accuracy, and fraction of 5’ UMI reads could be modulated as expected by varying the cDNA or Tn5 amounts (**Figure 2d,e**).

**Figure 2.**
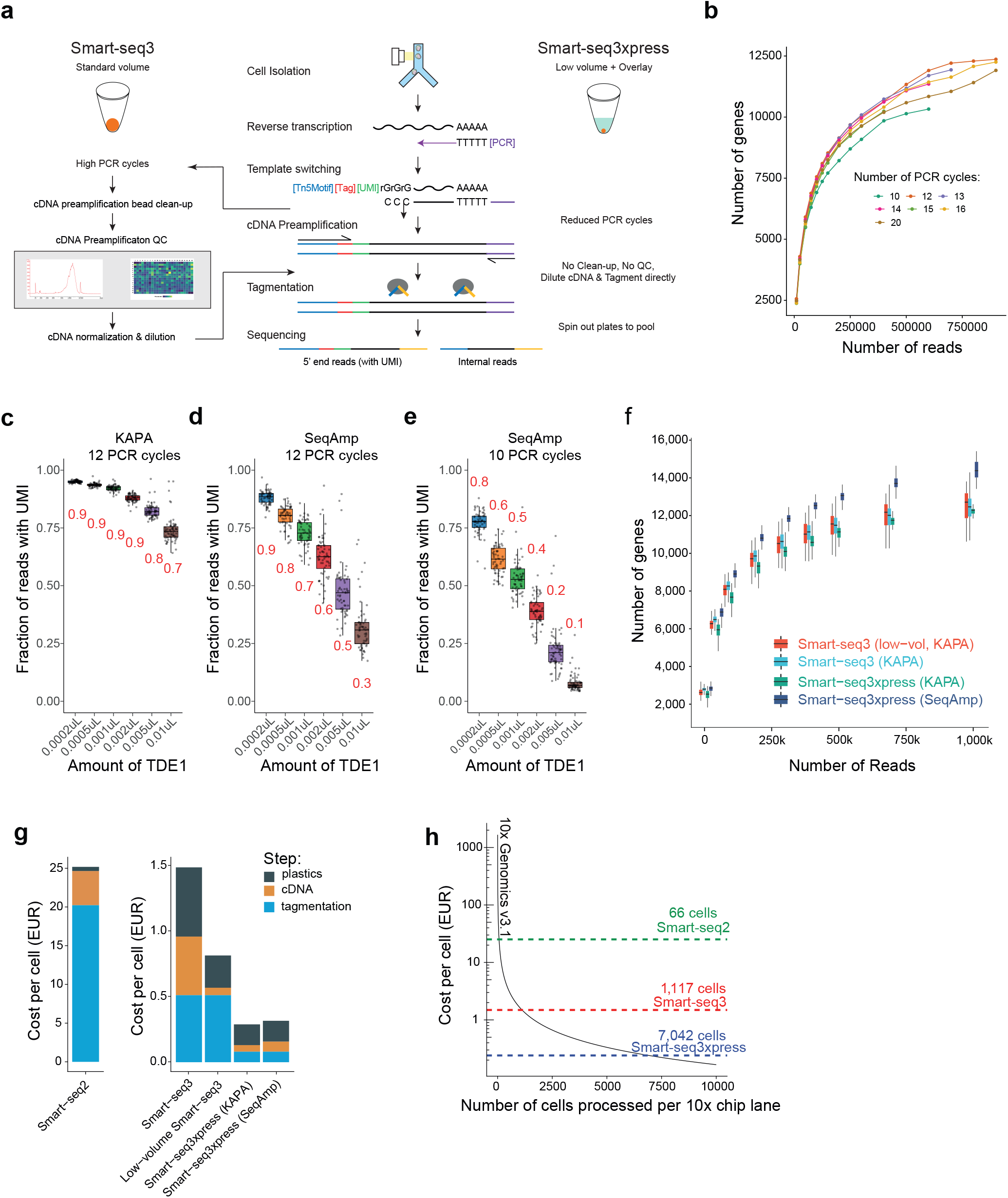
Scalable full-transcript coverage scRNA-seq with Smart-seq3xpress. (**a**) Schematic outline of the Smart-seq3 and Smartseq3xpress workflows. (**b**) Smart-seq3xpress is buffered for use with various amounts of pre-amplification PCR cycles. For each dot, the median number of Genes is calculated from the indicated number of raw sequencing reads in at least n>= 8 HEK293FT cells. (**c**) Fraction of UMI containing reads to internal reads for HEK293FT cells prepared with Smartseq3xpress (KAPA HiFi; 12 PCR cycles), at a variable range of TDE1 amounts. Boxplot shows median quartile range (n=64 each). (**d**) Fraction of UMI containing reads to internal reads for HEK293FT cells prepared with Smartseq3xpress (SeqAmp; 12 PCR cycles), at a variable range of TDE1 amounts. Boxplot shows median quartile range (n=60 each). (**e**) Fraction of UMI containing reads to internal reads for HEK293FT cells prepared with Smartseq3xpress (SeqAmp; 10 PCR cycles), at a variable range of TDE1 amounts. Boxplot shows median quartile range (n= 60, 60, 60, 62, 64, 64, respectively). (**f**) Benchmarking of Smart-seq3 variants. Shown are the number of genes detected in HEK293FT cells in the full-volume Smart-seq3 (Hagemann-Jensen et al., 2020), low-volume Smart-seq3 and Smart-seq3xpress implementations at the indicated read depths (n>= 4). (**g**) Cost of Smart-seq library preparations by summing up list prices of reagents. Shown is the cost per cell divided by costs relating to cDNA synthesis (Lysis, RT, PCR), tagmentation (including indexing PCR) and plastics. (**h**) Approximate cost per cell of Smart-seq library preparations and a single lane of 10x Genomics v3.1 relative to the number of cells analyzed.

Next, we benchmarked both low-volume Smart-seq3 (1 μL, KAPA), Smart-seq3xpress (12 PCR cycles, using either KAPA or SeqAmp), with the recently published data from the standard volume Smart-seq3^3^. Importantly, we observed better gene and molecule detection as a function of sequencing depth for SeqAmp-based Smart-seq3xpress (**Figure 2f** and **Supplementary Figure 8a**). Thus, the miniaturization of reaction conditions and omitting several experimental steps of Smart-seq3 was indeed feasible at improved gene and molecular detection and simultaneously making it feasible to reach sequencing-ready Smart-seq3xpress libraries within a single workday. Importantly, the cost per Smart-seq3xpress singe-cell library are reduced to 0.25 EUR (**Figure 2g**), similar in cost to processing ~7,000 cells on a 10x Genomics v3.1 3’ gene expression lane (**Figure 2h**). Further streamlining and reduction of plastics consumables was achieved by collecting final libraries by centrifugation using a simple 3D printed adapter (**Supplementary Figure 9**) and through contact-less combinatorial index dispensing, and relying on tagmentation plates containing already dispensed desiccated index primers.

We processed 16,349 human peripheral blood mononuclear cells (hPBMCs) with Smart-seq3xpress and sequenced the libraries on a DNBEQ G400RS (MGI) to an average depth of 230,000 read pairs per cell. Sequence data was demultiplexed, processed and quality controlled with zUMIs^12^ and Seurat^13^ was used to analyze data and project cells using UMAP (**Methods**). The single-cell transcriptomes separated into 23 clusters (**Figure 3a**) that were supported by all donors (**Figure 3c**) and corresponding to the main immune cell types expected in hPBMCs. Reconstruction of T-cell receptor sequences from the internal reads matched the identified T-cell clusters (**Figure 3b**), and our cluster assignments were in good agreement with cell type labels inferred by the reference-based integration in Azimuth (**Supplementary Figure 10)**. Recent studies showed that scRNA-seq alone was not capable of separating some of the known cell types and states in hPBMCs without the help of additional protein measurements. For example, a recent 10x Genomics based study^13^ sequenced 200,000 hPBMCs transcriptomes and required up to 228 protein markers (CITE-seq) to distinguish several cell types and states, including unconventional T-cell population (MAIT cells), gamma-delta T-cells and effector memory T-cells. Strikingly, all these cell types separating by the Smart-seq3xpress transcriptomes alone (**Figure 3a**), exemplified with marker gene expression for MAIT and gamma-delta T-cells (**Figure 3e**), and further validated by their specific T-cell receptors (data not shown). Moreover, we identified the separation of additional rare cell populations, including a CD4+ T-cell population only characterized by their clonal expression of specific T-cell receptors (**Figure 3d,e**). Notably, we obtained this unprecedented level of granularity in *de novo* cell type and state characterization using only 16,349 Smart-seq3xpress transcriptomes, compared to droplet-based analysis of RNA and proteins in 200,000 individual cells. Thus, to generate high-quality transcriptomes seem as important as obtaining high cell numbers in order to accurately characterize cell types and states in complex samples and tissues. Thus, we conclude that Smart-seq3xpress can generate unprecedented high-quality full-transcript coverage scRNA-seq data from complex human samples, at highly competitive cost and time.

**Figure 3.**
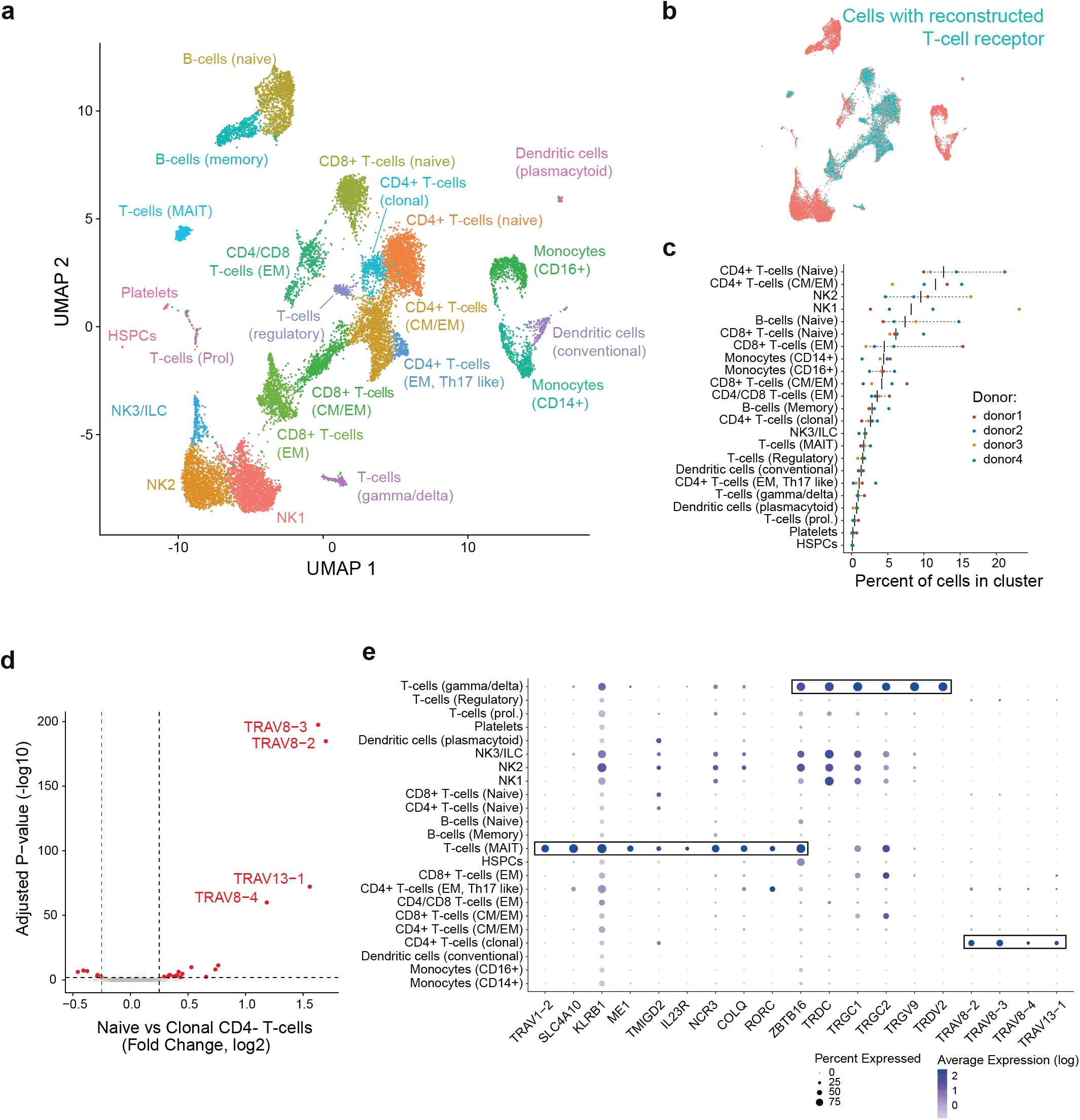
Smart-seq3xpress analysis of human PBMCs. (**a**). Dimensional reduction (UMAP) of 16,349 human PBMCs from 4 donors produced with Smartseq3xpress (KAPA) colored and annotated by cell type. (EM = Effector Memory), (CM = Central Memory), (NK = Natural Killer), (ILC = Innate Lymphoid Cells), (HSPC = Hematopoietic Stem and Progenitor Cell), (MAIT = Mucosal Associated Invariant T-cell). (**b**) Overlay onto UMAP of presence of detected of T-cell Receptor (TCR) using TRaCeR to reconstruct the TCR info from the sequenced Smartseq3xpress data. (**c**) Distribution of cell type abundances as a percentage of all cells from each of the 4 donors. (**d**) Differential gene expression analysis between Naïve CD4 T-cell cluster (n=2,064) and Clonal CD4 T-cell cluster (n=384). Indicated are the top four TCR genes driving the clonal CD4 T cell cluster separation. (**e**) Dotplot showing expression of selected marker genes for MAIT, gamma-delta and clonal CD4 T cells in all annotated clusters with size of the dot denoting the detection of a gene within the cells of the cluster and color scale denoting the average expression level.

Large-scale efforts to enumerate cell types and states across tissues in humans and model organisms (e.g. NIH Brain Initiative and Human Cell Atlas) are dominated by single-cell RNA-sequencing that count RNA expression at only the 5’ or 3’ end, since commercial droplet-based technologies or combinatorial indexing methods have high throughput and lower costs. With the development of Smart-seq3xpress, we demonstrate that it is possible to rival throughput and cost while capturing full-transcript information with nanoliter plate-based automation. Not only has Smart-seq3xpress overcome the main limitations of plate-based assays in terms of per-cell cost, but the data obtained also challenges the existing notions of the cell numbers required for efficient and finely resolved clustering of cells. Therefore, high sensitivity scRNA-seq with isoform- and allele-specific resolution can for the first time be performed with Smart-seq3xpress at a scale suitable for large-scale cell atlas building.

## Methods

### Cell sources, culturing and sorting

HEK293FT cells (Invitrogen) were grown in DMEM medium (4.5 g/l glucose, 6 mM L-glutamine, Gibco), supplemented with 10% fetal bovine serum (FBS) (Sigma-Aldrich), 0.1 mM MEM nonessential amino acids (Gibco), 1 mM sodium pyruvate (Gibco) and 100 μg/ml pencillin-streptomycin (Gibco) at 37 °C. For sort of single cells, cells were harvested by incubation with TrypLE Express (Gibco). K562 cells were grown in RPMI medium, supplemented 10% FBS, and 1% pencillin-streptomycin. Frozen aliquots of 10 million human PBMCs from healthy individuals were purchased from Lonza, requiring healthy donors only. Written informed consent was obtained at sampling point from all donors by Lonza and our analyses of hPBMCs were approved by the Regional Ethical Review Board in Stockholm, Sweden (2020-05070). hPBMCs were gently thawed before sorting. For all cell types, dead cells were gated out after staining with propidium iodide (Thermo Fisher). Live, single cells were sorted into 384-well plates containing lysis buffer using a BD FACSMelody equipped with a 100-μm nozzle and plate cooling. (BD Biosciences). After sorting each plate was immediately spun down and stored at −80 degrees.

### Smart-seq3 library preparation

Full-volume Smart-seq3 library preparations were performed as previously published^3^. Briefly, cells were sorted into 3 μL lysis buffer containing 0.1 % Triton-X100, 5% PEG8000 adjusted to 4uL RT volume, 0.04 μL RNAse Inhibitor (Takara; 40 U/μL), 0.5μM oligo-dT (IDT; 100uM 5′-biotin-ACGAGCATCAGCAGCATACGA-T30VN-3′) and 0.5mM dNTPs/each (Thermo Fisher; 25 mM each) and snap frozen at −80 °C. After denaturation for 10min at 72C, reverse transcription was initiated by addition of 1μL of RT mix, containing (25mM Tris-HCl~ pH 8.3 (Sigma), 30mM NaCl (Ambion), 0.5mM GTP (Thermo Fischer Scientific), 2.5mM MgCl2 (Ambion), 8mM DTT (Thermo Fischer Scientific), 0.25U/μL RNase Inhibitor (Takara), 2μM template switching oligo (TSO: 5’-Biotin-AGAGACAGATTGCGCAATGNNNNNNNNrGrGrG-3’; IDT), and 2U/μL Maxima H-minus Reverse transcriptase (Thermo Fischer Scientific). Reverse transcription was carried out at 42 degrees for 90min followed by 10 cycles of 50 degrees for 2min and 42 degrees for 2 min, terminating at 85C for 5min. 6μL PCR mix (1x KAPA HiFi PCR buffer(Roche), 0.3mM dNTPs/each(Roche), 0.5mM MgCl2 (Ambion), 0.5μM Smart-seq3 Fwd Primer (5’-TCGTCGGCAGCGTCAGATGTGTATAAGAGACAGATTGCGCAATG-3’ ; IDT), 0.1μM Smart-seq3 Rev primer (5’-ACGAGCATCAGCAGCATACGA-3’ ; IDT), and 0.02U/μL KAPA HiFi DNA polymerase (Roche)), is subsequently added to each well. PCR was carried out using following termal conditions: 3min at 98 degrees for initial denaturation, 20 cycles of 20 secs at 98 degrees, 30 sec at 65 degrees, 6 min at 72 degrees.

### Low volume Smart-seq3 & Smart-seq3xpress library preparation

For experiments including overlays, including Vaporlock (Qiagen), Silicon Oils (Sigma), Tri-/Tetradecane (Sigma), 3μL of designated overlay was added to each well, and stored at room temperature until use. Lysis buffer of various volumes (0.1μL – 3μL) were dispensed using a either Formulatrix Mantis, or Dispendix I-Dot one liquid dispenser to each well, all containing (0.1% TX-100, 5% PEG8000 adjusted to RT volume, 0.5μM Oligo-dT adjusted to RT volume, 0.5mM dNTPs/each adjusted to RT volume, 0.5U RNase Inhibitor (Takara; 40 U/μL). After dispensing, lysis plates were briefly centrifuged down to ensure lysis is properly collected and stored under the overlay. Stored plates of sorted cells were denatured at 72 °C for 10min, followed by addition of indicated volumes of reverse transcription mix, common for all is the reagent concentrations are stable; 25mM Tris-HCl~ pH 8.3 (Sigma), 30mM NaCl (Ambion), 0.5 mM GTP (Thermo Fisher Scientific), 2.5mM MgCl2 (Ambion), 8mM DTT (Thermo Fischer Scientific), 0.25U/μL RNase Inhibitor (Takara), 2uM template switching oligo (TSO: 5’-Biotin-AGAGACAGATTGCGCAATGNNNNNNNNrGrGrG-3’; IDT), and 2U/μL Maxima H-minus Reverse transcriptase (Thermo Fischer Scientific). After RT mix dispensing the plate was spun down to ensure merge of RT and lysis reactions. Reverse transcription was performed at 42 degrees for 90min followed by 10 cycles of 50 degrees for 2min and 42 degrees for 2 min. Indicated volumes of PCR master mix was dispensed, all containing constant reaction concentrations of (1x KAPA HiFi PCR buffer(Roche), 0.3mM dNTPs/each(Roche), 0.5mM MgCl2 (Ambion), 0.5μM Smart-seq3 Fwd Primer (5’-TCGTCGGCAGCGTCAGATGTGTATAAGAGACAGATTGCGCAATG-3’ ; IDT), 0.1μM Smart-seq3 Rev primer (5’-ACGAGCATCAGCAGCATACGA-3’ ; IDT), and 0.02U/μl KAPA HiFi DNA polymerase (Roche). After dispensing plate was quickly spun down before incubated in PCR as follows: 3min at 98 degrees for initial denaturation, 10-24 cycles of 20 secs at 98 degrees, 30 sec at 65 degrees, 2-6 min at 72 degrees. Final elongation was performed for 5 min at 72 degrees. For conditions with after cDNA preamplification clean-up; 100nl of either H2O, ExoSAP-IT express and 0.5U ExoI + 0.05 FAST-AP was dispensed per well and incubated at 37 degrees for 15min followed by inactivation at 85 degrees for 5 min.

For Smartseq3xpress with SeqAmp (Takara), lysis and RT was carried out with 2μM TSO unless otherwise indicated as described above. PCR mastermix was dispensed at 0.6μL per cell containing (1x SeqAmp PCR buffer, 0.025U/μL SeqAmp polymerase, 0.5μM/1μM Smartseq3 Fwd and Rev Primer). After dispensing PCR mastermix the plate was quickly spun down before incubated as follows: 1min at 95 degrees for initial denaturation, 6-18 cycles of 10 secs at 98 degrees, 30 sec at 65 degrees, 2-6 min at 68 degrees. Final elongation was performed for 10 min at 72 degrees. For Smartseq3xpress with NEBNext Ultra II Q5 Master Mix (NEB), PCR mastermix consisted of (1x NEBNext Ultra II Q5 Master Mix, 0.5μM/1μM Smartseq3 Fwd and Rev Primer) and was performed at 30sec at 98 degrees for initial denaturation, 12 cycles of 10 secs at 98 degrees, 30 sec at 65 degrees, 6 min at 72 degrees. Final elongation was performed for 5 min at 72 degrees. For Smartseq3xpress with NEBNext Q5 Hot Start HiFi PCR Master Mix (NEB), PCR mastermix consisted of (1x NEBNext Q5 Hot Start HiFi PCR Master Mix, 0.5μM/1μM Smartseq3 Fwd and Rev Primer) and was performed at 30sec at 98 degrees for initial denaturation, 12 cycles of 10 secs at 98 degrees, 30 sec at 65 degrees, 1 min at 65 degrees. Final elongation was performed for 5 min at 65 degrees. For Smartseq3xpress with Platinum SuperFi II DNA polymerase (Invitrogen), PCR mastermix consisted of (1x SuperFi II Master Mix, 0.2μM dNTPs, 0.5μM/1μM Smartseq3 Fwd and Rev Primer) and was performed at 30 sec at 98 degrees for initial denaturation, 12 cycles of 10 sec at 98 degrees, 30 sec at 60 degrees, 6 min at 72 degrees. Final elongation was performed for 5 min at 72 degrees. For Smartseq3xpress with Platinum II Taq Hot Start DNA polymerase (Invitrogen), PCR mastermix consisted of (1x Platinum II Taq Master Mix, 0.2μM dNTPs, 0.5μM/1μM Smartseq3 Fwd and Rev Primer) and was performed at 2min at 94 degrees for initial denaturation, 12 cycles of 15 secs at 94 degrees, 30 sec at 60 degrees, 6 min at 68 degrees. Final elongation was performed for 5 min at 68 degrees.

### After preamplification workflow

For regular Smart-seq3, preamplified cDNA libraries were purified with home-made 22% PEG beads at a ratio of 1:0.6. Library sizes were observed utilizing Agilent Bioanalyzer High Sensitivity Chip, followed by concentration quantification using QuantiFlour dsDNA assay (Promega). cDNA was subsequently diluted to 100pg/μL, unless otherwise specified.

For low volume preamplified cDNA libraries were diluted by addition of 9μL H2O to each well, if not indicated otherwise, followed by a quick centrifugation. Library sizes were checked on an Agilent Bioanalyzer, using the High sensitivity DNA chip, meanwhile concentrations were quantified using QuantiFlour dsDNA assay (Promega). cDNA was normalized to 100pg/μL if nothing else specified.

For Smart-seq3xpress preamplified cDNA libraries were diluted with the addition of 9μL H2O unless stated otherwise, before transferring 1μL of diluted cDNA from each well into tagmentation.

### Sequence library preparation for Smart-seq3xpress

Tagmentation was performed in 2 μL consisting of 1μL of either diluted or normalized preamplified cDNA and 1μL of 1x tagmentation buffer (10mM Tris pH 7.5, 5mM MgCl2, 5% DMF), (0.025-0.5) μL ATM (Illumina XT DNA sample preparation kit) or (0,0002-0,01) μL TDE1 (Illumina DNA sample preparation kit). In the event of in-house Tn5 1x tagmentation buffer used consisted of (10mM TAPS-NaOH pH 8.4, 5mM MgCl2, 8% PEG8000) and indicated amounts of Tn5 0.0005-0.01 μM inhouse TN5 enzyme. Samples were incubated at 55 degrees for 10min, followed by addition of 0.5μL of 0.2% SDS to each well. After addition of 1.5μL custom Nextera index primers carrying 10bp dual indexes, library amplification was started by the addition of 4μL PCR mix (1x Phusion Buffer (Thermo Scientific), 0.01 U/μL Phusion DNA polymerase (Thermo Scientific), 0.2 mM dNTP/each) and incubated at 3 min 72 degrees; 30 sec 95 degrees; 12-14 cycles of (10 sec 95 degrees; 30 sec 55 degrees; 30 sec 72 degrees); 5 min 72 degrees in a thermal cycler. Samples were pool by spinning out each plate gently in a 300mL robotic reservoir (Nalgene) fitted with a custom 3D printed scaffold by pulsing to ~200g. The pooled library was purified with home-made 22% PEG beads, at a ratio of 1:0.7.

### Sequencing

Libraries were sequenced on a NextSeq500 or DNBSEQ G400RS platform. For NextSeq runs, denatured libraries were loaded on HighOutput v2.5 cartridges at 2.1 - 2.3 pM. For G400RS runs, libraries were created using phosphorylated index primers or subjected to 5 cycles of adapter conversion PCR using the MGIEasy Universal Library Conversion Kit (MGI) and subsequently circularised from 1 pmol of dsDNA according to the manufacturer’s protocol. 60 fmol of circular ssDNA library pools were used for DNA nanoball (DNB) making using a custom rolling-circle amplification primer (5‘-TCGCCGTATCATTCAAGCAGAAGACG-3’). DNBs were loaded on FCL flow cells (MGI) and sequenced PE100 using custom sequencing primers (Read 1: 5’-TCGTCGGCAGCGTCAGATGTGTATAAGAGACAG-3’; MDA: 5’-CGTATGCCGTCTTCTGCTTGAATGATACGGCGAC-3’, Read 2: 5’-GTCTCGTGGGCTCGGAGATGTGTATAAGAGACAG-3’; i7 index: 5’-CCGTATCATTCAAGCAGAAGACGGCATACGAGAT-’3;CTGTCTCTTATACACATCTGACGCTGCCGACGA-’3). i5 index: 5’-CTGTCTCTTATACACATCTGACGCTGCCGACGA-’3).

### Primary data processing

zUMIs^12^ version 2.8.2 or newer was used to process raw fastq files. Reads were filtered for low quality barcodes and UMIs (4 bases < phred 20, 3 bases < phred 20 respectively) and UMI-containing reads parsed by detection of the pattern (ATTGCGCAATG). Reads were mapped to the human genome (hg38) using STAR version 2.7.3 and error-corrected UMI counts were calculated from Ensembl gene annotations (GRCh38.95). zUMIs was also used to downsample cells to equal raw sequencing depth to facilitate method benchmarking.

### Analysis of hPBMC data

Cells were filtered for low quality libraries, requiring i) more than 50% of read pairs mapped to exons+Introns, ii) more than 20.000 read pairs sequenced, iii) more than 500 genes (exon+intron quantification) detected per cell, and less than 15% of read pairs mapped to mitochrondrial genes. Furthermore, a gene was required to be expressed in at least 10 cells. Analysis was done using Seurat v4.0.1^13^. Data was normalized (“LogNormalize”), scaled to 10,000, and total number of counts, and mitochondrial fraction was regressed out. Utilizing the Seurat integration function, the donor effect, from the 4 different donors in the dataset was removed. The top 10,000 variable genes were considered and 35 principal components for SNN, neighborhood construction and UMAP dimensionality reduction. Cell clusters were produced using Louvain algorithm at a resolution of 0.8. Cell types were identified by using either Seurat FindAllmarkers (Wilcoxon), or the R package Presto (Wilcoxon & AUC). For the Azimuth predictions, a qc filtered count matrix was uploaded to the Azimuth web-based application, and processed according to the Azimuth app.

### TCR reconstruction

T cell receptor sequences were reconstructed using TraCeR v0.6.0^14^ run in the teichlab/tracer Docker environment and using the --loci A B G D --species Hsap flags.

### Molecular Spike data processing & analysis

Molecular spike11 data was extracted from aligned zUMIs bam files and analysed using the UMIcountR package (https://github.com/cziegenhain/UMIcountR; v0.1.1). After loading the data using the “extract_spike_dat” function, overrepresented spikes were discarded with a read cutoff of 25 and above. We next used molecular spike observations across all cells conditions with at least 5 reads per molecule to sample 26 ground truth mean expression levels from 1 to 316 molecules per cell using the “subsample_recompute” function. We then plotted the mean counting difference shaded by the standard deviation.

## Acknowledgements

We are grateful to Gert-Jan Hendriks for designing and 3D printing the adapter for library pooling. We thank the Sandberg Lab for fruitful discussions. This work was supported by grants to R.S. from the Swedish Research Council, the Knut and Alice Wallenberg Foundation, and the Göran Gustafsson Foundation. C.Z. was supported by an EMBO long-term fellowship (ALTF 673–2017).

## Data and code availability

All sequencing data will be deposited under ArrayExpress accession (pending) at the European Bioinformatics Institute (EBI). Processing of Smart-seq3 and Smart-seq3xpress libraries was performed in zUMIs (https://github.com/sdparekh/zUMIs) and additional information can be made available upon request.

## Author Contributions

Developed and optimized the method: MHJ. Designed the experimental infrastructure for miniaturization and automation: CZ, Performed experiments and analysed data: CZ, MHJ. Conceived study and wrote manuscript: CZ, MHJ, RS. Supervised work: RS.

## Supplementary Figure Legends

**Supplementary Figure 1:**
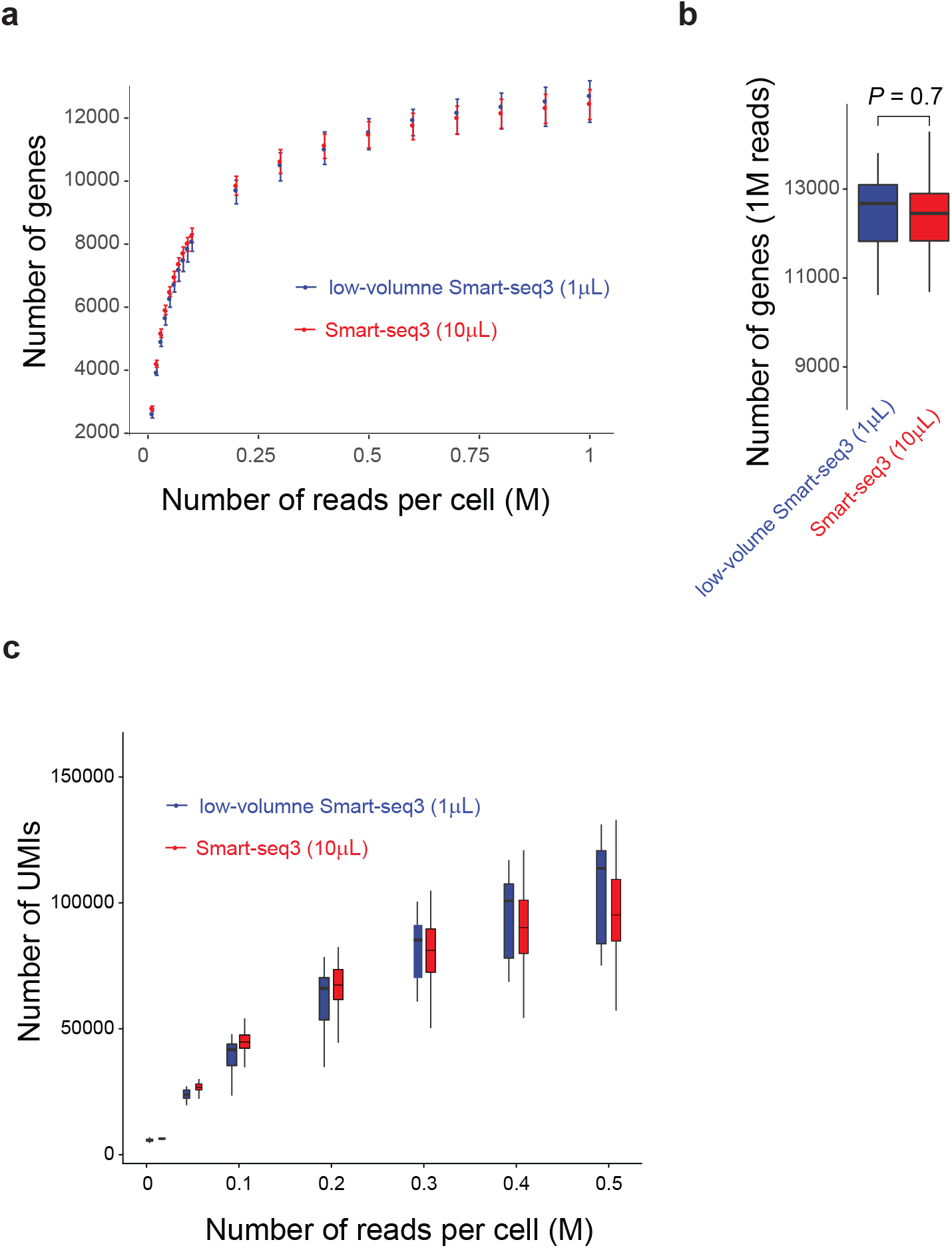
Benchmarking of low-volume cDNA synthesis. (**a**) Number of genes detected in low-volume Smart-seq3 and published Smart-seq3 data from HEK293FT cells. Each point denotes the median number of genes at each sequencing depth with the lines indicating upper and lower quartiles (n >= 8 cells each). (**b**) Number of genes detected at 1 million reads per cell. Indicated is the p-value of a two-sided t-test (p=0.7). (**c**) Number of UMIs detected in low-volume Smart-seq3 and published Smart-seq3 data from HEK293FT cells. Each point denotes the median number of UMIs at each sequencing depth with the lines indicating upper and lower quartiles (n >= 8 cells each).

**Supplementary Figure 2:**
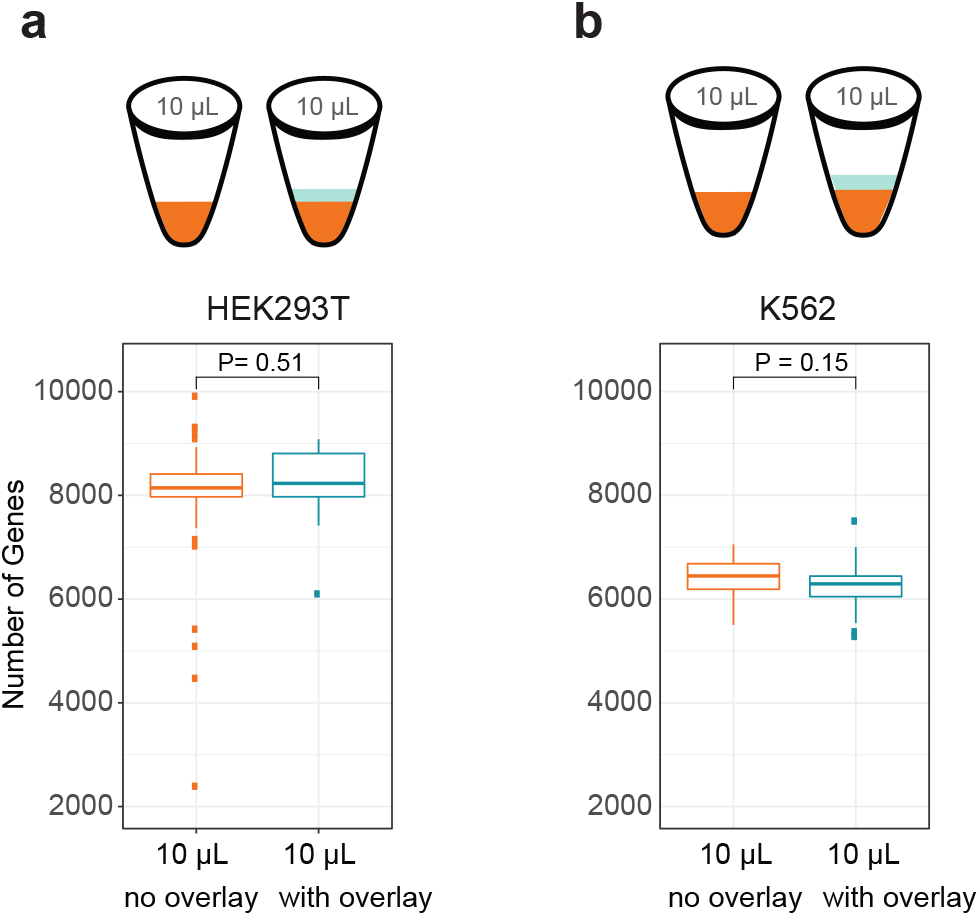
Influence of Vapor-lock overlay on standard Smart-seq3. For both (**a**) HEK293FT and (**b**) K562 cells the influence of creating standard volume Smart-seq3 libraries without (HEK293FT n=37; K562 n=26) and with an overlay (VaporLock; HEK293FT n=15; K562 n=31)) was compared. Boxplots show genes detected and p-values show result of a two-sided t-test.

**Supplementary Figure 3:**
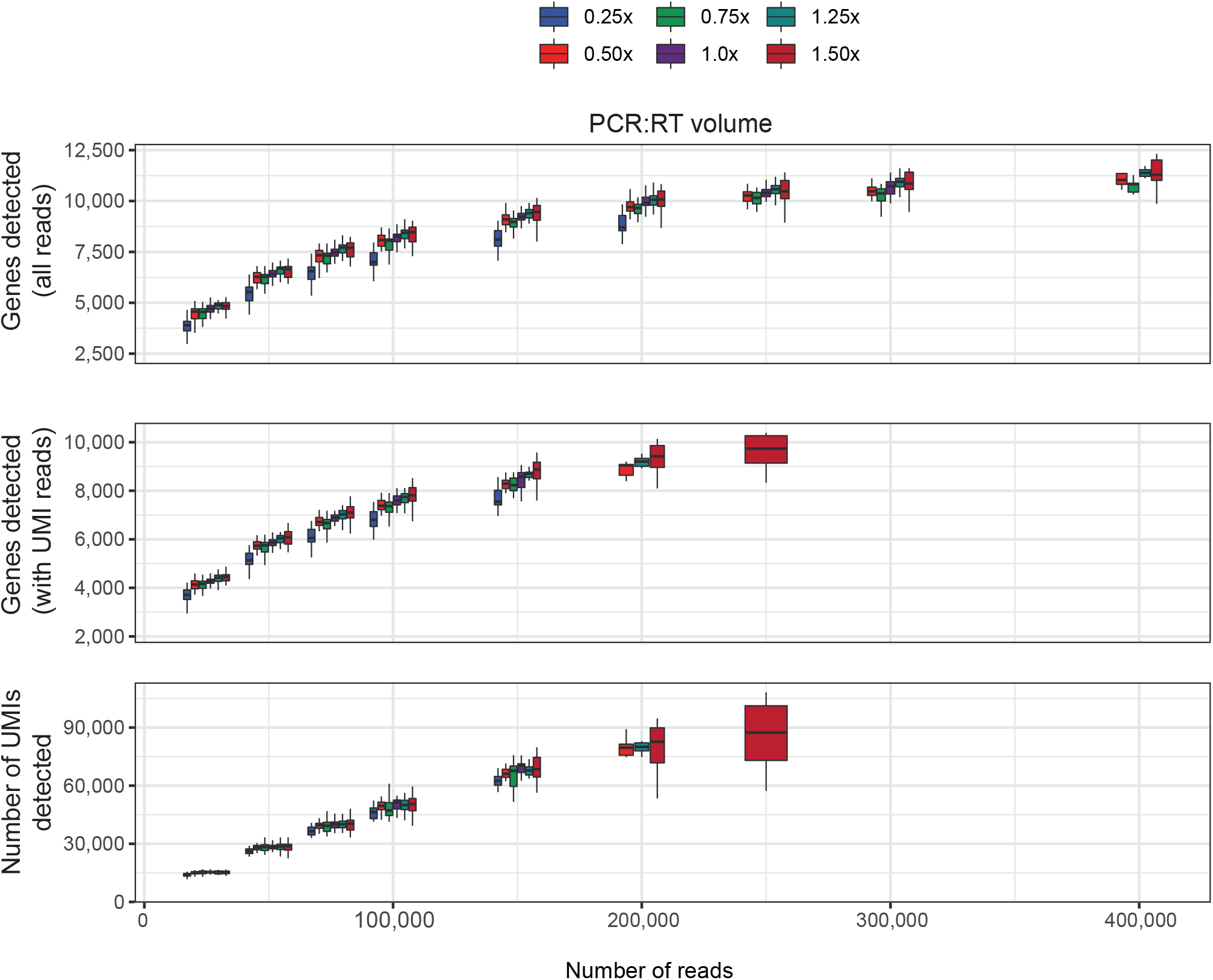
Ratio of PCR volume to RT volume. For indicated ratios, boxplots show genes detected, genes with UMIs detected and UMIs captured, downsampled by sequenced reads. At each depth and condition, boxplots (n>=5) indicate the median, upper and lower quartiles of the observed complexity)

**Supplementary Figure 4:**
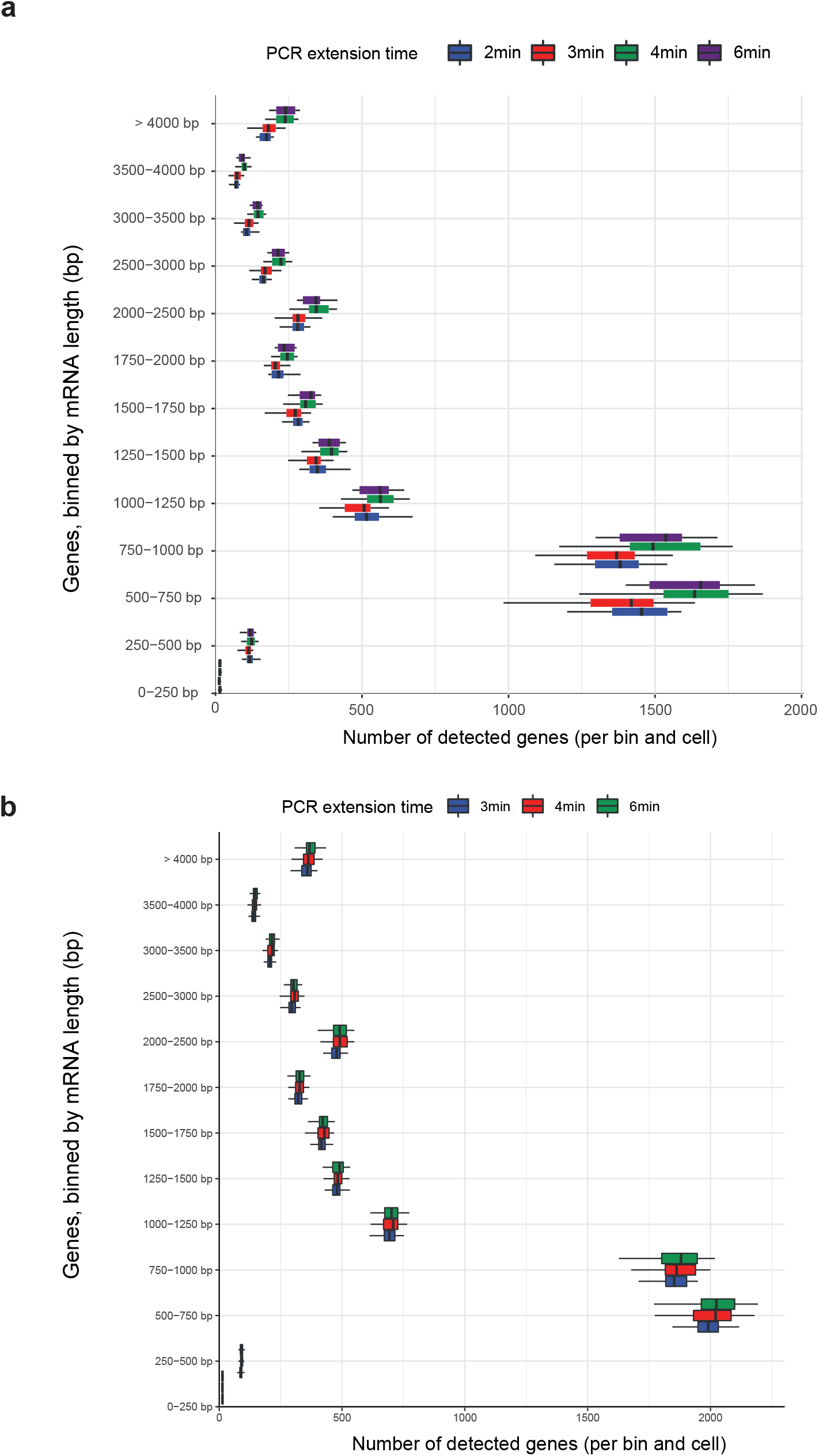
Influence of PCR extension time for KAPA HiFi Hot Start and SeqAmp polymerases. (**a**) Boxplots for each extension time (2, 3, 4, 6 min) using KAPA HiFi Hot Start polymerase are shown as detected genes binned by their transcript length at 250,000 reads per cell. (n=29, 24, 29, 10; respectively) (**b**) Number of genes detected binned by transcript length for each extension time 3 min (n = 64), 4min (n = 64), 6min (n = 60) using SeqAmp polymerase at 250,000 reads per cell.

**Supplementary Figure 5:**
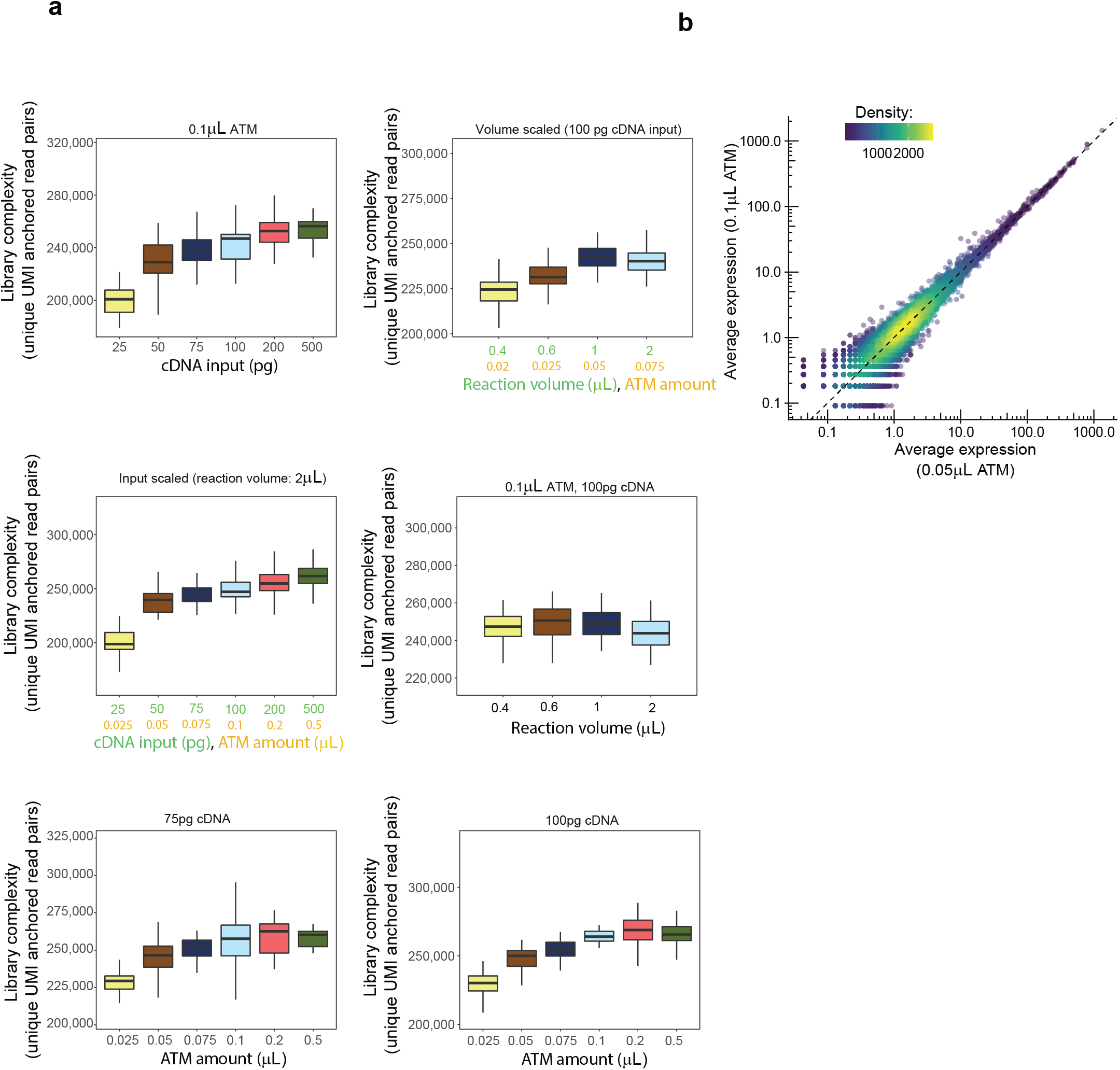
Investigating tagmentation complexity. (**a**) Systematic investigation of tagmentation complexity was performed by varying cDNA input with constant Tn5 amount, varying input and Tn5 amounts, varying Tn5 amounts with constant cDNA input, scaling of reaction volumes and Tn5 amounts with constant cDNA input, scaling of reaction volumes with constant cDNA and Tn5 amounts. For each boxplot, shown is the library complexity in terms of unique gene-assigned UMI-anchored read pairs (unique per-molecule cut-sites) from 400,000 raw sequencing reads. Each condition contains between 22 and 73 HEK293FT cells. (**b**) Concordance of gene expression levels between HEK293FT cells tagmented using 0.05 μL ATM Tn5 and 0.1 μL ATM Tn5 (mean UMI counts).

**Supplementary Figure 6:**
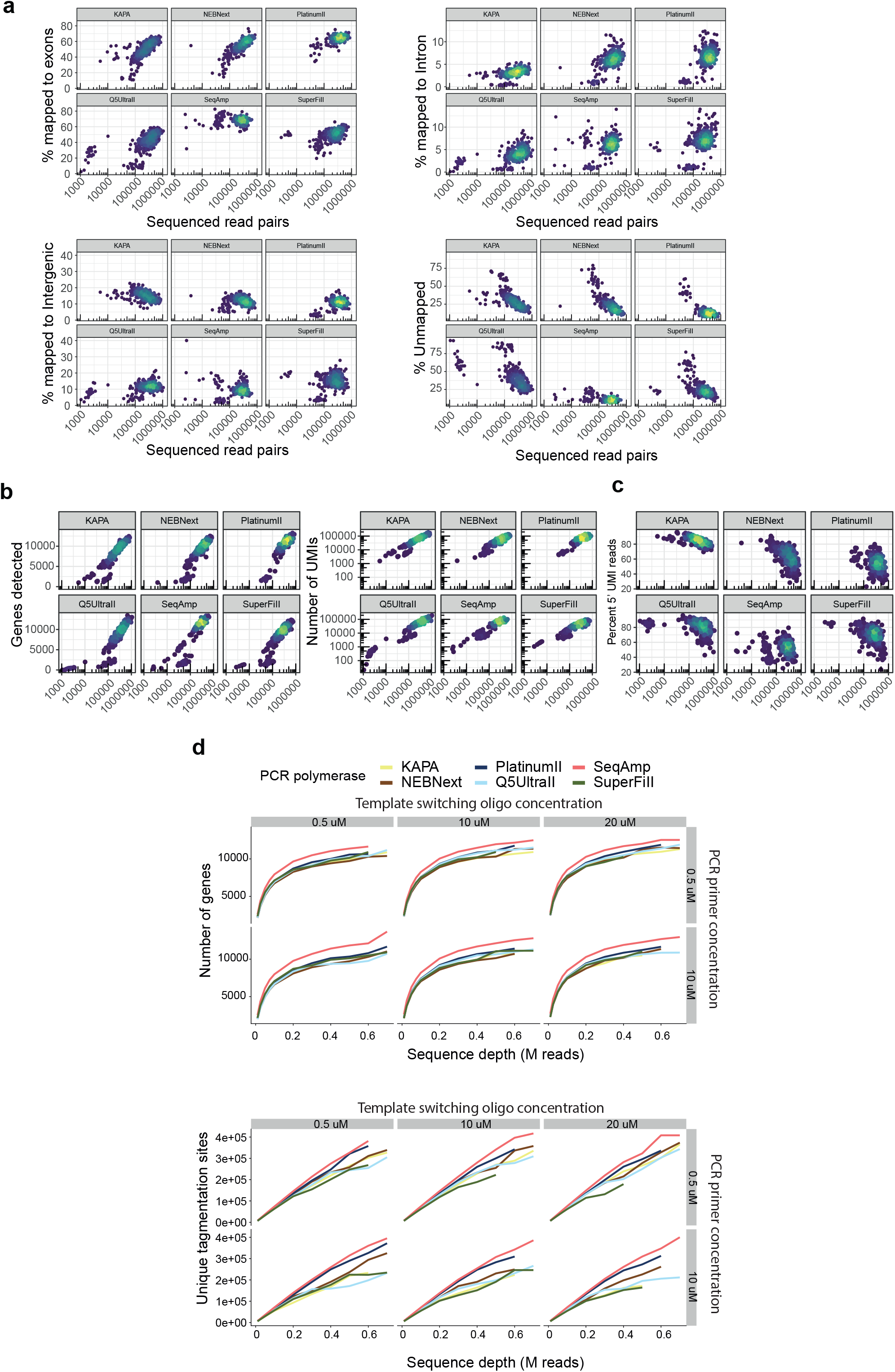
Performance of pre-amplification polymerases for Smartseq3xpress. (**a**) Mapping statistics of the six different tested commercial polymerases: KAPA HiFi Hot Start (n=384), NEBNext Q5(“NEBNext”, n=384), NEBNext Ultra II Q5(“Q5UltraII”, n=384), Platinum II (n=384), SuperFi II (n=384), SeqAmp (n=360). Dotplots show percentage of mapped reads to exons, introns, intergenic regions and unmapped with the sequenced read depth. (**b**) Number of Genes and UMIs detected per cell relative to sequencing depth for all six polymerases tested. (**c**) Fraction of UMI containing reads versus internal reads for each of the tested polymerases. (**d**) Lines show median number of genes detected and median number of unique tagmentation sites downsampled by sequenced reads using 0.5 μM, 1 μM and 2 μM TSO concentration in RT, combined with either 0.5 μM or 1μM of eachfForward and reverse PCR primer (n >= 56 cells per condition).

**Supplementary Figure 7:**
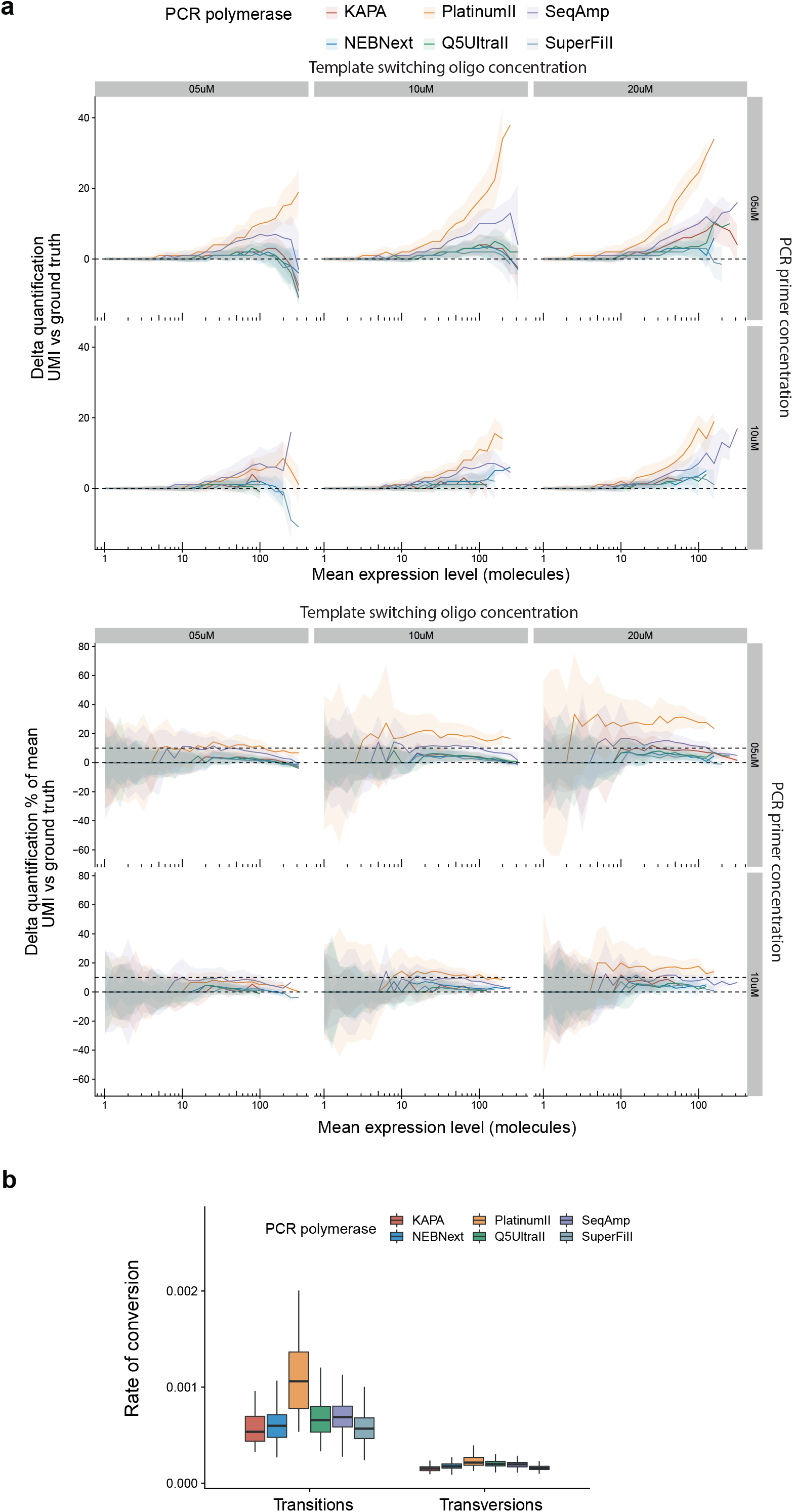
Counting accuracy and error-rate of PCR polymerases. (**a**) Molecular spike-ins were used to assess the accuracy of each polymerase in mRNA molecule counting, based on the counting difference between internal molecular spikes counts and Smart-seq3xpress UMIs at indicated amounts of TSO and PCR primer concentrations. Colored lines indicate the mean counting difference between the unique spike identifiers and quantified UMIs when sampling the spike at the indicated mean expression levels for each of the polymerases, shaded by the standard deviation. The counting differences are expressed in absolute deviance or relative to the mean molecule number. (**b**) Rate of base conversions in aligned reads relative to the reference genome. For every polymerase, we compute the average fraction of transitions and transversions, shown as boxplots over all cells (n >= 360 in each condition).

**Supplementary Figure 8:**
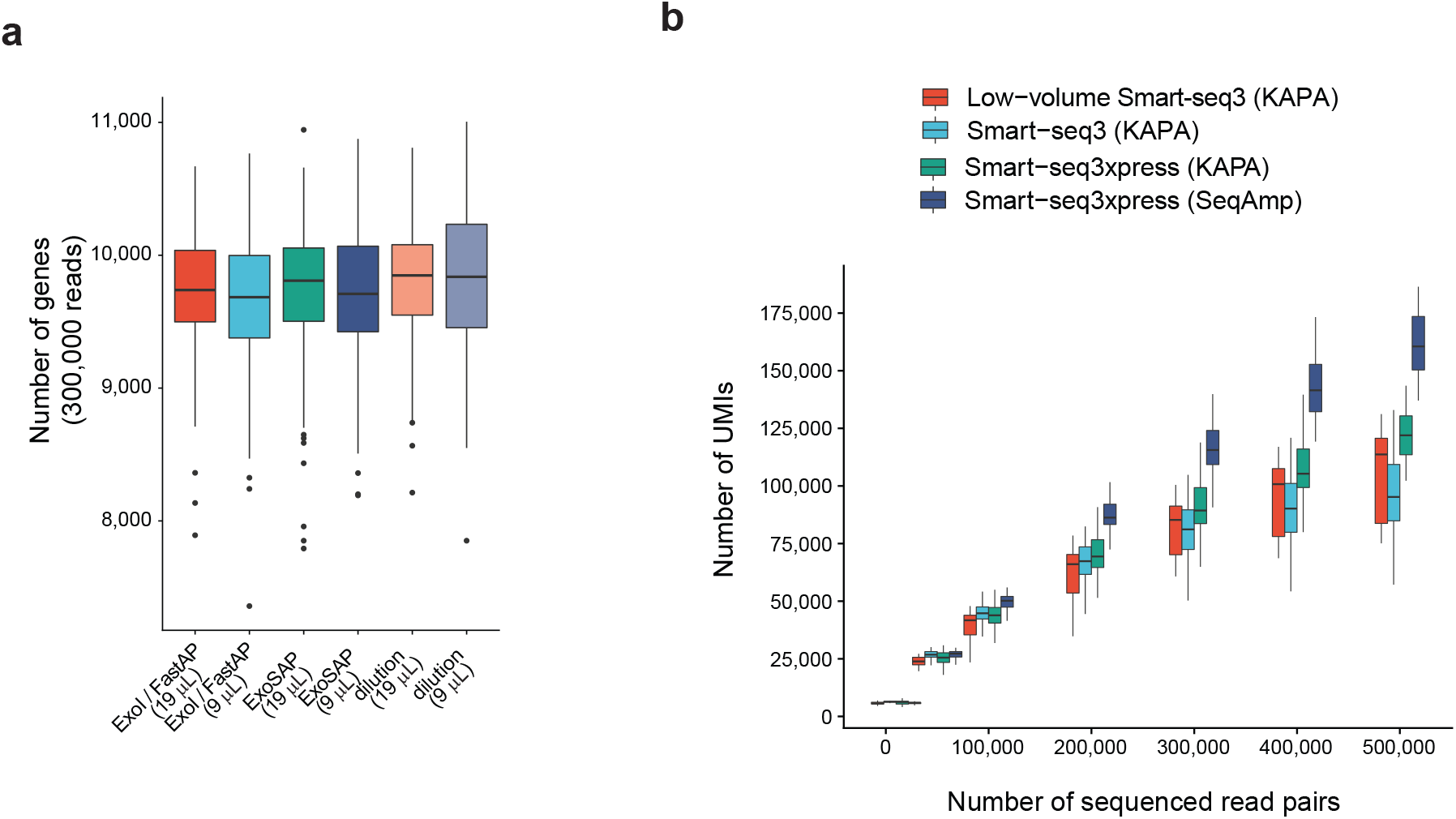
Influence of post PCR clean up and UMI capture benchmarking for Smartseq3xpress. (a) Genes detected after treating preamplified cDNA libraries to reduce “contaminants”, such as dNTPs and oligonucleotides, before going into tamentation. All conditions (dilution alone, ExoSAP IT-express, or ExoI + Fast-AP (ExoSap)) were either diluted in 9 or 19uL of water. (b) Benchmark of number of UMIs detected in HEK293FT cells in the full-volume Smart-seq3 (Hagemann-Jensen et al., 2020), low-volume Smart-seq3 and Smart-seq3xpress implementations using KAPA and SeqAmp at the indicated read depths (n>= 4).

**Supplementary Figure 9:**
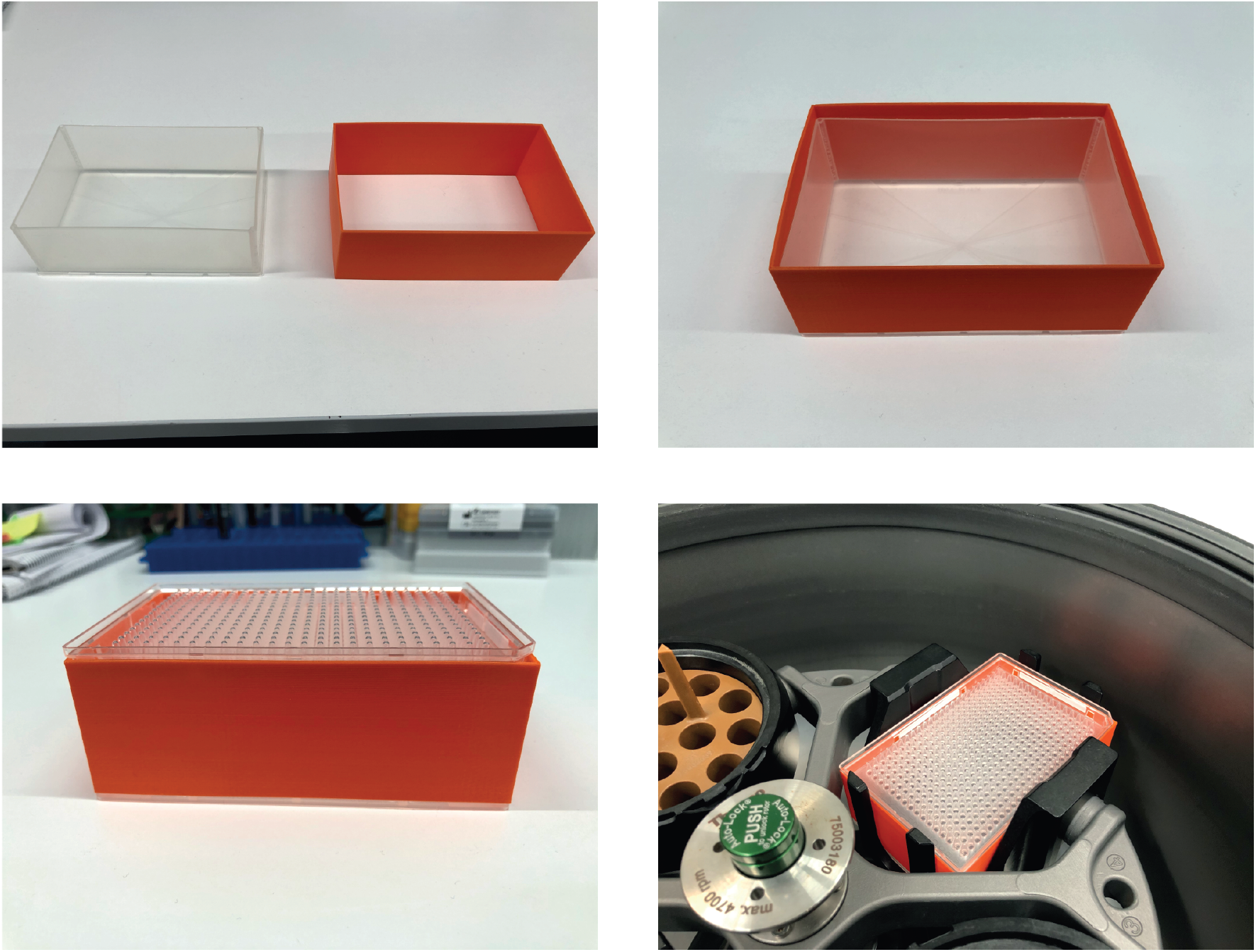
3D printed adapter to facilitate PCR plate pooling by centrifugation. Pictures showing reservoir and 3D printed frame/holder and assembly to facilitate pooling of 384 well plates by quick and gentle centrifugation.

**Supplementary Figure 10:**
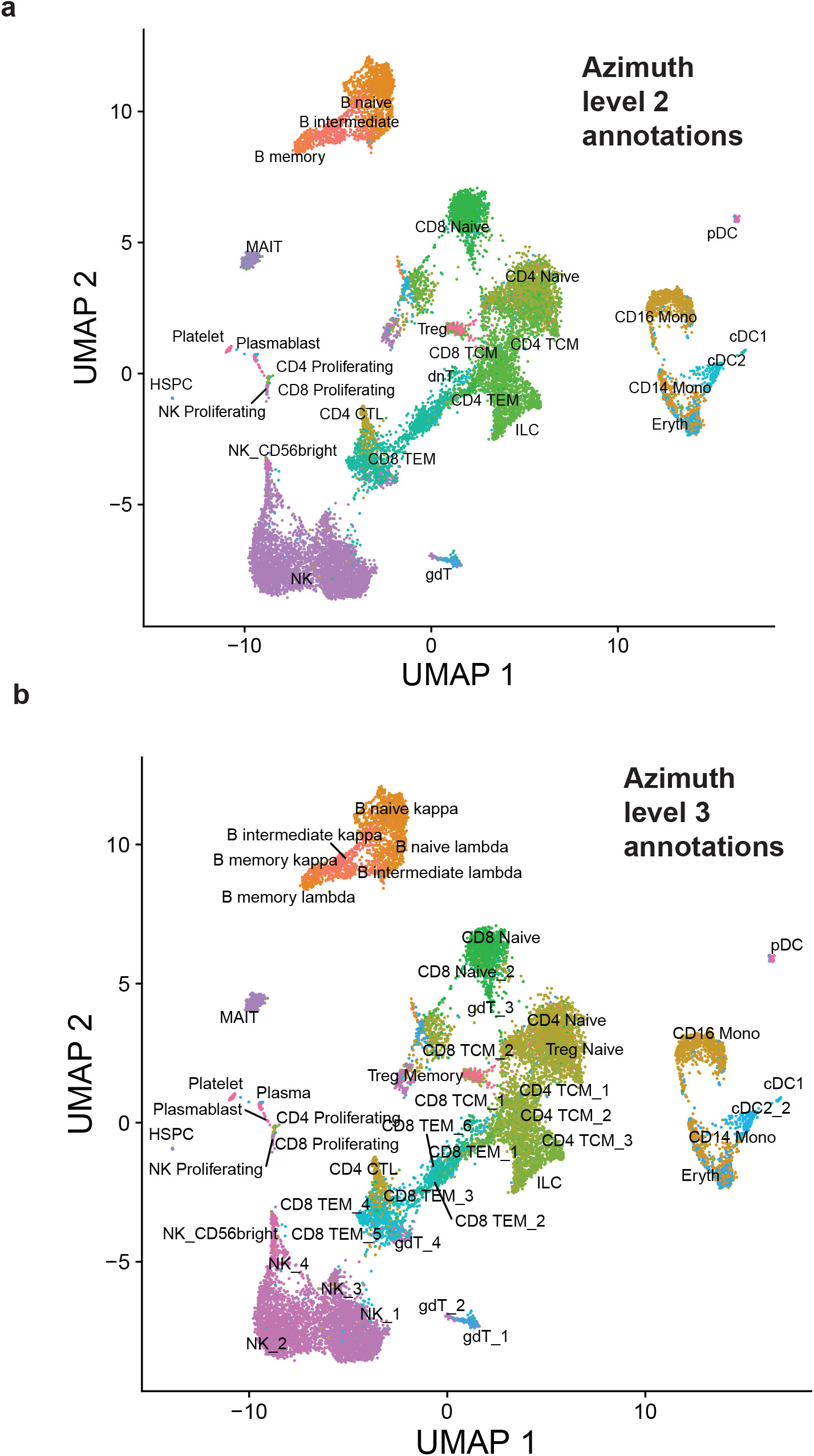
Smartseq3xpress hPBMCs annotated with Azimuth. Smartseq3xpress hPBMC generated in this study was annotated by reference-based prediction and annotation from Azimuth (Hao et al., 2020) using the hPBMC reference dataset available at https://azimuth.hubmapconsortium.org. (**a**) Smartseq3xpress UMAP (as in Figure 3a) is colored and annotated by Azimuth cell type annotation level 2. (**b**) Smartseq3xpress UMAP is colored and annotated by Azimuth cell type annotation level 3.

## References

1. Ziegenhain, C. et al. Comparative Analysis of Single-Cell RNA Sequencing Methods. Mol Cell 65, 631–643.e4 (2017).

2. Mereu, E. et al. Benchmarking single-cell RNA-sequencing protocols for cell atlas projects. Nature Biotechnology (2020) doi:10.1038/s41587-020-0469-4.

3. Hagemann-Jensen, M. et al. Single-cell RNA counting at allele and isoform resolution using Smart-seq3. Nature Biotechnology 38, 708–714 (2020).

4. Muraro, M. J. et al. A Single-Cell Transcriptome Atlas of the Human Pancreas. Cell Syst 3, 385–394.e3 (2016).

5. Tabula Muris Consortium et al. Single-cell transcriptomics of 20 mouse organs creates a Tabula Muris. Nature 562, 367–372 (2018).

6. Mayday, M. Y., Khan, L. M., Chow, E. D., Zinter, M. S. & DeRisi, J. L. Miniaturization and optimization of 384-well compatible RNA sequencing library preparation. PLoS One 14, e0206194 (2019).

7. Mamanova, L. et al. High-throughput full-length single-cell RNA-seq automation. Nat Protoc 16, 2886–2915 (2021).

8. Mora-Castilla, S. et al. Miniaturization Technologies for Efficient Single-Cell Library Preparation for Next-Generation Sequencing. J Lab Autom 21, 557–567 (2016).

9. Jaeger, B. N. et al. Miniaturization of Smart-seq2 for Single-Cell and Single-Nucleus RNA Sequencing. STAR Protoc 1, 100081 (2020).

10. Picelli, S. et al. Tn5 transposase and tagmentation procedures for massively scaled sequencing projects. Genome Res 24, 2033–2040 (2014).

11. Ziegenhain, C., Hendriks, G.-J., Hagemann-Jensen, M. & Sandberg, R. Molecular spikes: a gold standard for single-cell RNA counting. bioRxiv 2021/451877, (2021).

12. Parekh, S., Ziegenhain, C., Vieth, B., Enard, W. & Hellmann, I. zUMIs - A fast and flexible pipeline to process RNA sequencing data with UMIs. Gigascience 7, (2018).

13. Hao, Y. et al. Integrated analysis of multimodal single-cell data. Cell 184, 3573–3587.e29 (2021).

14. Stubbington, M. J. T. et al. T cell fate and clonality inference from single-cell transcriptomes. Nat Methods 13, 329–332 (2016).

